# Natural spider silk enhances mechanical performance of collagen scaffold under stretching conditions

**DOI:** 10.1101/2025.03.27.645864

**Authors:** Maryia Tsiareshyna, Sammi Y.T. Huang, Chen-Pan Liao, Ming-Jer Tang, Tzyy Yue Wong, I-Min Tso

## Abstract

Collagen is the most abundant protein in the extracellular matrix, crucial for wound healing and cell proliferation. While it holds promise as a scaffold for tendon, skin, and ligament reconstruction, collagen’s mechanical strength, particularly under stretch, is poor. Previous attempts to improve collagen strength involved blending it with silkworm or recombinant spider silk. In this study, for the first time, we evaluated whether collagen gel from fish skin could be strengthened by infusing it with native spider silk, specifically the major ampullate (MA) silk of *Nephila pilipes*, known for superior mechanical properties. MA silk was woven onto a frame, pressed into a PDMS platform, and then used to create a collagen scaffold. Young’s modulus of the infused collagen scaffold, subjected to either stretching or non-stretching treatments, was measured using AFM. After 24 hours of cyclic stretching, collagen infused with silk showed less fragility, higher Young’s modulus, and no bacterial growth. Immunohistochemical staining showed that after stretching, the thickness and architecture of the collagen gel infused with silk were maintained, and the fibers were reorganized in a more compact, aligned, and denser manner. Overall, collagen infused with native spider silk exhibited improved mechanical stability and stiffness under cyclic stretching, suggesting that this combination could serve as a robust matrix for bioengineering applications while preventing bacterial infiltration.

## Introduction

Collagen is the most abundant protein in the extracellular matrix and plays a role in healing wounds and enhancing cell proliferation and adhesion. Collagen also has a great potential as scaffold for tendon, skin and ligament reconstruction. The interaction between collagen and β-1 integrin can regulate cell adhesion and proliferation [1,2,3,4,5]. Type I collagen can be used to produce autografts to support bone defect repair in orthopedics. It provides osteoinductive and pro-angiogenic molecular signals, osteoconductive matrix support and mechanical stability [6]. Collagen-based hydrogels are biocompatible, biodegradable, hemostatic, have adjustable physicochemical properties and demonstrate both thermal and enzymatic stability. Properties such as mechanical stability can also be modified or improved by special additives such as polyethylene glycol and polyvinyl alcohol. Physical, chemical or enzymatic crosslinking strategies have been used to modify collagen hydrogel properties for specific therapeutic purposes such as controlling inflammation or inhibiting bacteria in wound healing [7]. Collagen is also utilized in the recovery of tendon and ligament injuries. To create bio-tendon constructs, collagen gels were combined with human-induced pluripotent stem cells (iPSCs), and cyclic mechanical stretching with progressively increasing loads was applied for two weeks. Transplantation of such bio-tendon construct exhibited an aligned extracellular matrix structure with good cell infiltration properties [8].

Despite employing various crosslinking strategies, it remains challenging to enhance the mechanical properties and control the degradation rate of colla^3–5^gen. Silk is a biopolymer known for its remarkable extensibility and strength, attributed to its glycine-rich and alanine-rich motifs. Collagen scaffold combined with knitted *Bombyx mori* silk was utilized for the repair of the medial collateral ligament (MCL) in rabbits. [2]. Since internal spaces were created to allow neoligament growth so tissue regeneration was enhanced. Collagen matrix incorporated with knitted silk not only had better mechanical properties, but it also facilitated interface healing between the neoligament and the native ligament. The fibril diameter of collagen + silk-treated MCLs was smaller than that of native ligament; however, it was still larger than silk-treated MCLs and exhibited greater mechanical properties. Connective tissues have more space to form in knitted than braided or parallel fiber scaffolds, which is necessary for ligament repair. Implanted collagen combined with *B. mori* silk scaffolds in mice over three months demonstrated biocompatibility and maintained 25.6% of their original tensile strength [2]. Panaz-Perez et al.[3] reported that collagen scaffolds integrated with *B. mori* silk fibers exhibited tensile strength comparable to, or even exceeding, that of the human anterior cruciate ligament. Additionally, Zheng et al. [4] explored the use of collagen scaffolds embedded with *B. mori* silk for treating rotator cuff tears. The inclusion of *B. mori* silk enhanced the porosity, inductivity, and mechanical strength of the 3D microporous collagen scaffold. This design supported tendon cell migration, and the integration of tendon cells with collagen fibers resulted in a structure resembling natural tendon, indicating significant potential for shoulder injury regeneration. The study also reinforced previous findings, showing that a higher collagen concentration increases structural stability by reducing porosity and promoting cell infiltration.

However, one limitation of the model proposed by Zheng et al. [4] was the high collagen degradation rate. Kwon et al. [9] reported that implanting collagen scaffold incorporated with *B. mori* silks could enhance regeneration of tendon tissue, which tensile properties and diameter were similar to those of native tendon. Concentration of silk fibers (SF) in scaffolds was shown to play a significant role in influencing microstructure and morphology. High concentration of silk fibers enhanced formation of SF microfibers and microspheres in collagen scaffold which could effectively guide cell infiltration and improve hemocompatibility [10]. Collagen scaffolds implanted with heparin-loaded silk fibers could promote neovascularization and therefore can potentially be applied in vascular graft. For corneal tissue engineering, Long et al. [11] created collagen scaffolds with different ratio of silk fibroin and used cross-linking facilitating 1-ethyl-3-(3-dimethylaminopropyl) carbodiimide. They found that a 100:10 collagen and silk fibroin ratio could generate optimal mechanical properties and high suture retention strength. Additionally, membrane made with such ratio was suitable for proliferation of human corneal epithelial cells and complete epithelialization was achieved in 35 ± 5 days [12]. For cartilage tissue regeneration in osteoarthritis treatment, Lin et al. [13] created scaffolds composed of different ratio of type II collagen and silk fibroins. They found that a 70:30 collagen-silk fibroin ratio composite membrane exhibited the best mechanical stability, cell proliferation ability, swelling ratio and degradation rate. Therefore, collagen plus silk fiber composite made from such combination has the most promising potential for cartilage engineering. Liu et al. [14] evaluated the effects of collagen fiber alignment and silk fiber incorporation on tendon and bone regeneration. They found that after eight weeks of implantation in rabbits more regenerative tissues were found on randomly-aligned collagen scaffolds incorporated with silk fibers.

Spider dragline silk is recognized for its superior mechanical properties compared to silkworm silk, prompting some researchers to attempt improving the mechanical properties of collagen by incorporating this material. However, because spiders only produce a small quantity of silks and silks from certain taxa with good mechanical properties are difficult to obtain, some researchers used recombinant silks instead. Zhu et al. employed the electrospinning technique to fabricate aligned fibers using collagen and spider dragline proteins sourced from transgenic goats.. Recombinant silk proteins could increase stability of mimicked native extracellular matrix (ECM) and improve properties such as deformation resistance and fiber swelling. Moreover, the mechanical properties of collagen scaffold could be enhanced by increasing silk protein concentration. However, the tensile strength, stiffness and stretchability of recombinant spider silks produced by electrospinning were considerably lower than those of natural spider silks due to imperfection of molecular assembly during this process. Pure collagen fibers exhibited a mean ultimate tensile strength of 40 MPa, an elastic modulus of 0.58 GPa, and an ultimate strain of 12.1%. In contrast, natural spider major ampullate silk fibers had corresponding values of 182 MPa for ultimate tensile strength, 4.45 GPa for elastic modulus, and 5.6% for ultimate strain [15]. The tensile strength, elasticity, fiber stability and stretchability could be improved on a short term basis by treating the collagen constructs with glutaraldehyde vapor, which increased β sheets in silks and linked collagen peptides [5]. Another challenge of applying collagen in cell culture is the undesired swelling of such hydrogels due to hydration of hydrophilic amino acids while being incubated in medium. Researchers found that by increasing quantity of silks in collagen scaffold the water content could be decreased and consequently the swelling could be controlled. On the other hand, the quantity of collagen in scaffold plays a vital role in tissue engineering because cell adhesion and expression of β-1 integrin, a receptor necessary for intracellular signaling, performed better when collagen content is high. Currently, researchers are using electrospinning to create collagen-silk protein composite exhibiting optimal fiber density and matrix stiffness to facilitate cell proliferation and differentiation.

Cells *in vivo* experience various mechanical stimulations such as contraction, relaxation, blood flow, and stiffness of the ECM. Dynamic microenvironment characterizes how cells interact with ECM and such condition is necessary for the survival of cells. Cyclic stretching under different magnitudes, frequencies, and periods of stretch ally collagen fiber bundles together. There is evidence showing that stretching can reduce the size of breast tumor in mice by 52% [16]. Stretching also triggers apoptosis of cancer cells by influencing cell migration. During the initial stretching cycle, cancer cells change their structure and size uniquely, which phenomenon might be applied in early cancer diagnosis [17]. Tijore et al. [18] showed that it was possible to selectively kill cancer cells by inducing mechanoptosis while stimulating growth of normal cells.

When subjected to cyclic stretching at a frequency of 1.5 Hz, collagen gel constructs cultured in the presence of elastogenic factors (EFs) demonstrated peak elastin mRNA expression and the highest accumulation of total matrix elastin [19]. Furthermore, magnetic stretching was applied to human periodontal ligament stem cells (hPDLSCs) within an aligned rat collagen scaffold, providing significant evidence of extracellular matrix (ECM) remodeling, a critical process in connective tissue development [20]. Stretching can enhance initial ECM degradation and new ECM secretion [21]. Collagen gel with sphere loaded with growth factor (GF) increased smooth muscles cell (SMC) morphology and protein expression after uniaxial cyclic stretch, which was important for successful SMC regeneration [22]. In another study, mature murine myogenic cell line C2C12 cells were cultured in collagen gel and cyclic stretch was applied. Such stimulation allowed mature myotubes to orient and form. This 3D reconstruction system for skeletal muscles has potential applications in the restoration of damaged muscles resulting from myopathy, injury, or the resection of malignant tumors [23].

In summary, although collagen-based gels exhibit wide variety of biomedical applications, one major limitation is its low mechanical performance and high degradation rate. It has been shown that recombinant spider silk proteins could significantly improve collagen gel matrix stability, strength, and cell adhesion. One of the novel approaches in many biomedical researches is cyclic stretching of tissues, which can also be applied to collagen gel constructs. However, although mechanical stimulation could enhance cell growth and protein production, collagen constructs cannot remain intact when subjected to such treatment. Stiffness of collagen gels severely decreased after stretching and therefore it could not replace animal models yet. Study on neurite extension showed that it was higher with the lowest collagen concentration in gel (0.6–0.8 mg/ml). Opposite to this mechanical stiffness of gels increased with increased concentration of collagen [24]. In this present study, we aim to create a strong collagen-based construct that exhibits desired stiffness after mechanical stimulation so that it could optimally support cell growth. Here, for the first time, we evaluated whether the mechanical properties of collagen gel retrieved from fish skin could be enhanced by infusing it with native spider silks. Although considerable efforts had been invested in artificial spider silks, the mechanical performances of recombinant silks were still inferior to those of natural spider silks [25]. The multicomponent nature of dragline silk, particularly its inclusion of MaSp1 and MaSp2 fibroins, has been instrumental in achieving its remarkable mechanical properties. Such structure has been a challenge for biomaterial research on artificial production of spider silks [26]. Natural spider silks also have surface layers such as lipid layer, glycoprotein layer, skin and core protein [27,28]. These layers may carry pheromones, pigments and antimicrobial agents [29,30,31] and might play roles in cell adhesion or aligning collagen fibers during stretching.

## Methods and materials

### Collagen gel construct making and stretching treatments

Giant wood spiders *Nephila pilipes* of the family Nephilidae are widely distributed in low and mid-elevation mountainous areas in Taiwan. Spiders were collected from Wushiken, Taichung City, Taiwan and were individually housed in enclosures at Tunghai University. We followed the method reported in Blamires et al. [32] to collect bundles of MA silks manually directly from large females through forcible extraction at a reeling rate of 0.01 m/sec. We irregularly wove (double sheets) native silk directly extracted from giant wood spider *N. pilipes* on a 2.8 × 2.8 cm plastic frame. Single spider silk sample contained 500 turns (approximately 10 mg of silk). Next, we placed frame with the silk into PDMS platform with sponge hooks on the sides. We pressed silk onto hooks and cut off the frame. Following this procedure, we made collagen gel by mixing: 0.26 ml of premixed buffer (5.7xDMEM, 2.5% NaHCO_3_, 0.1M HEPES, 0.17M CaCl_2_), 0.184 ml of medium, 0.543 ml of collagen (3.68 mg/ml density) and 1N NaOH. We made two types of collagen construct (with or without spider silk) and performed the following four treatments. In the first one, the collagen plus silk construct received cyclic stretching. In the second one, the collagen plus silk construct was not stretched. In the third one, the construct composed of collagen alone received cyclic stretching. In the fourth one, the construct composed of collagen alone was not stretched. The cyclic stretching of various collagen constructs was performed at 10% strain at a frequency of 1 Hz for a total of 24 hours. The collagen construct stretching was conducted using an ARTEMIS ATMS Boxer (TAIHOYA Corporation, Kaohsiung, Taiwan). During stretching the ATMS Boxer was placed in a CO_2_ incubator at a temperature of 37°C. The stretching device was composed of chambers which accommodated collagen gels on stretchable PDMS membranes (Genemessenger, Kaohsiung, Taiwan). The stretch adaptors were autoclaved with ozone before use. After cyclic stretching, the collagen gel were mounted on pre-prepared PDMS platforms placed in 3.5 cm petri dishes and were submitted to AFM measurements.

### AFM measurements

Measurement of cellular stiffness was taken according to method described previously [33]. Cantilever used was Arrow-TL1(Nanoworld, Neuchâtel, Swiss) modified with 5 μm (in diameter) polystyrene bead, spring constants were calibrated via thermal noise method in 1×PBS solution prior to each measurement valued at 0.02 to 0.08 N/m with indenting force of 1 nN. The rates of approaching and retracting for the cantilevel was 1 μm/sec. Force-distance curves were collected and calculated using JPK BioAFM software based on the Hertz model. A minimum of 50 cells were analyzed and data are presented as mean ± SEM.

### Statistical analyses

To compare the Young’s modulus among treatments, we built a Bayesian linear mixed model, which included sample identity and experiment trial nested within sample as nested random factor. These random factors allowed us to address the issue of non-random sampling. Fixed factors included stretching (stretched or not stretched), silk treatment (collagen only or collagen plus silk) and their interaction. Young’s modulus was transformed using the natural logarithm beforehand to normalize the sample distributions. We allowed the model to separately estimate the error variations for each treatment to cope with heteroscedasticity among treatments. We assigned the Student T distributions with three-degree freedom for the priors of differences between treatments and the 2.5 times scaled positive Student T distributions with three-degree freedom for the random terms. We reconstructed the posterior distributions of each parameter using 50,000 Markov chain Monte Carlo (MCMC) samples after a burn-in of 50,000 iterations. After model fitting, we summarized the posterior median and the 95% equal-tailed credible intervals of fixed effect. We reported the probability of direction (PD) and the Bayes factor (BF; estimated by using the Savage-Dickey density ratio) for fixed effect to infer the existence of an effect and the evidence ratio supporting the effect, respectively. All analyses were computed by using the R package “brms” version 2.21.0 [34] under the R framework version 4.4.1 [35].

### Immunohistochemistry staining

We used Masson’s Trichrome staining method to examine constructs composed of collagen gel alone and those infused with silk fibers. Samples were deparaffinized, hydrated through 100% alcohol, 95% alcohol 70% alcohol, and washed in distilled water. Next, samples were treated with Mordant dye in Bouins’s solution which had been preheated to 58 ° C for 15 minutes followed by rinsing in running water to remove the yellow color. Then the samples were stained in modified Weigert’s iron hematoxylin for 5 minutes and differentiated in acid-alcohol solution for 3-5 seconds. Next, stained in Biebrich Scarlet-Acid Fuchsin solution for 5 minutes, 1% Phosphomolybdic acid solution for 2 minutes, Aniline Blue solution for 5 minutes and acetic acid solution for 30 seconds. We rinsed our samples with 3-4 changes of distilled water after each of solution used. In the last step samples were dehydrated quickly through 95% ethyl alcohol and cleared using xylene.

## Results

### Effects of cyclic stretching on various gel constructs

Young’s modulus of collagen constructs receiving different treatments measured by AFM were given in Figure 1. After being cyclicly stretched for 24 hours, compared to gel constructs composed of collagen alone those infused with spider silks were less fragile, remained intact and had no bacteria growing on it. Additionally, gel construct composed of collagen and silk did not shrink and their Young’s modulus were significantly higher than those of constructs composed of collagen alone. Although the effect of interaction between amount of silk used in the gel construct and stretching was uncertain (PD = 0.932, BF = 0.99; Table 1), the results of post-hoc comparisons confirmed that the stretched samples composed of collagen and spider silk exhibited a Young’s modulus that was significantly twice as high as that of samples without spider silk (PDs > 98.6%, BFs > 5; Figure 2) and marginally significantly higher than non-stretched samples (PD = 97.7%, BF = 2.91; Figure 2).

**Figure 1.**
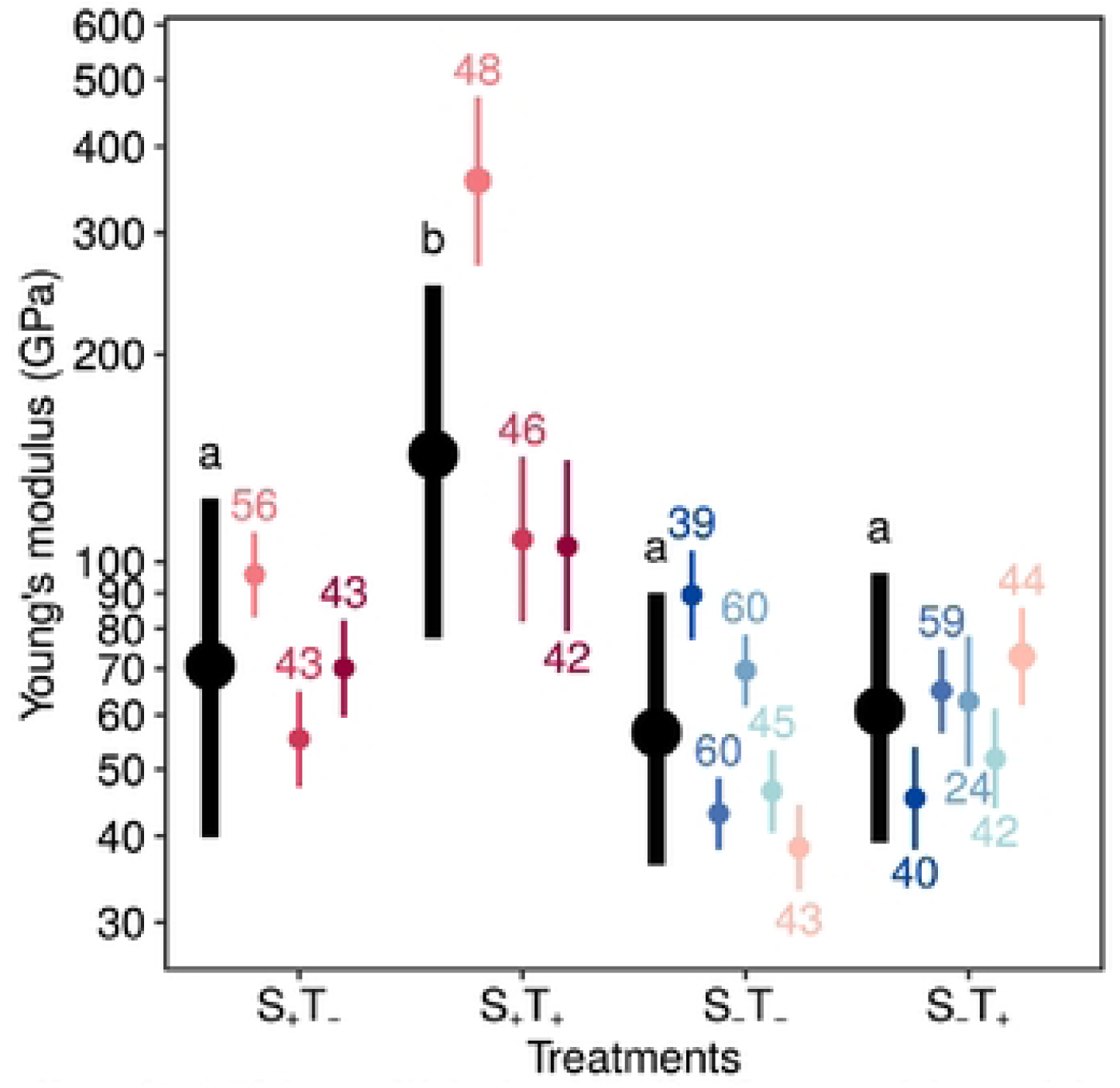
Posterior median(± 95% credible interval) of natural logarithm transformed Young’s modulus of collagen constructs receiving different treatments measured by AFM. Circles and error bars represent the posterior medians and 95% credible intervals respectively. The thick black bars represent the averages of each treatment. The colored thin bars represent the averages of each sample, and those bars in same color indicate the same sample. The number above or below the thin bars represents the number of tensile tests. Lowercase letters summarized the results of multiple comparisons based on the probability of direction. Abbreviations: S_+_, sample with spider silk; S_−_, sample without spider silk; T_+_, stretched sample; T_−_, unstretched sample.

**Figure 2.**
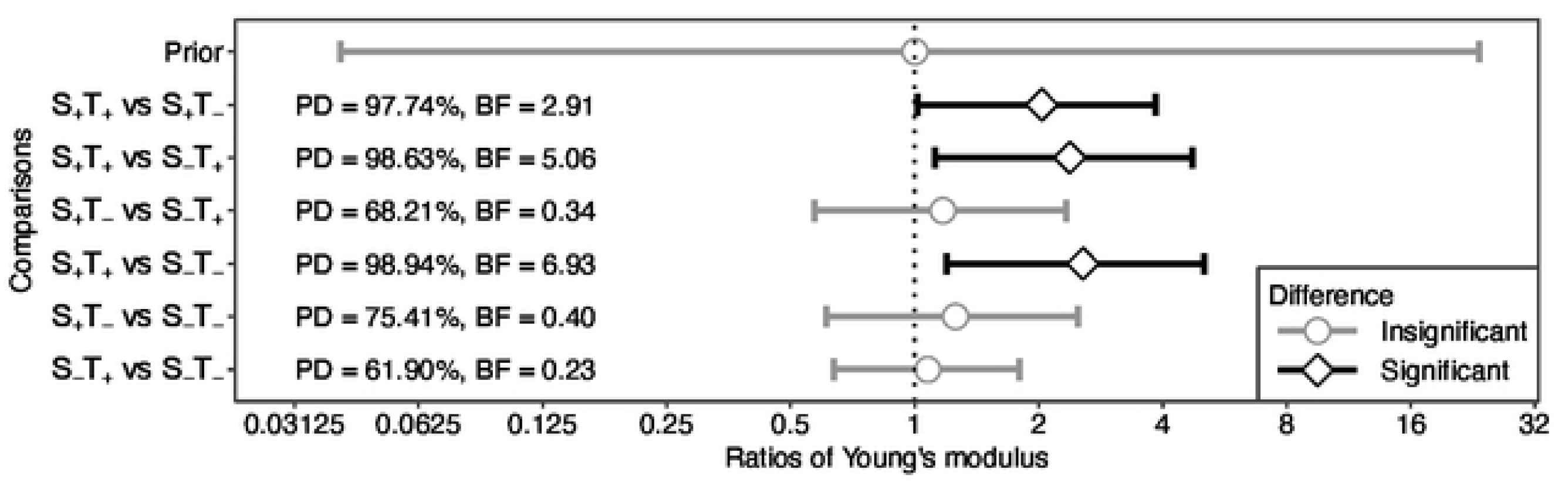
Summary of multiple comparisons in ratios of Young’s modulus among four treatments. Circles and error bars represent the posterior medians and 95% credible intervals (CrI), respectively. A ratio of Young’s modulus with a probability of direction (PD) greater than 97.5%, meaning the 95% credible interval (CrI) did not include 1, was denoted as significant. A Bayes factors (BF) greater than 3.16 indicates that the evidence supporting a difference is at least substantial. Abbreviations: S_+_, sample with spider silk; S_−_, sample without spider silk; T_+_, stretched sample; T_−_, unstretched sample.

**Table 1.**
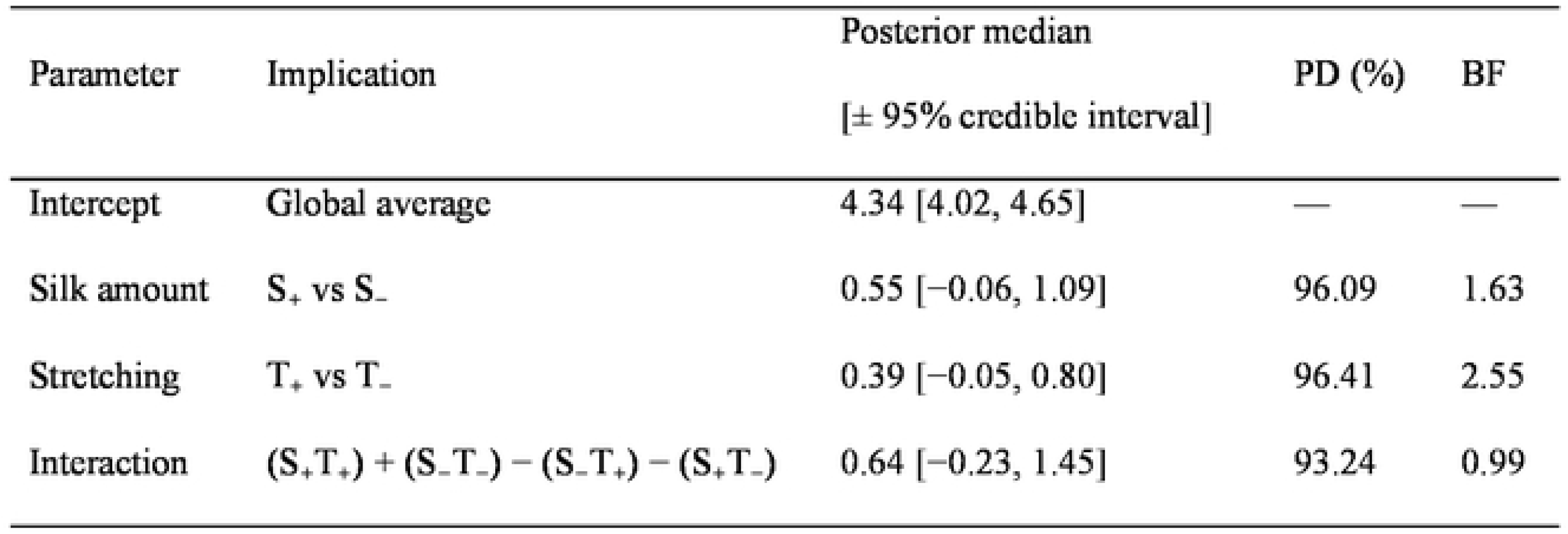
Summary of fixed effects in the model fitting natural logarithm transformed modulus. Abbreviations: S_+_, sample with silk; S_−_, sample without silk; T_+_, stretched sample; T_−_, non-stretched sample; PD, probability of direction; BF, Bayes factor.

| Parameter | Implication | Posterior median<br>[± 95% credible interval] | PD (%) | BF |
| --- | --- | --- | --- | --- |
| Intercept | Global average | 4.34 [4.02, 4.65] | — | — |
| Silk amount | S<sub>+</sub> vs S<sub>-</sub> | 0.55 [-0.06, 1.09] | 96.09 | 1.63 |
| Stretching | T<sub>+</sub> vs T<sub>-</sub> | 0.39 [-0.05, 0.80] | 96.41 | 2.55 |
| Interaction | (S<sub>+</sub>T<sub>+</sub>) + (S<sub>-</sub>T<sub>-</sub>) - (S<sub>-</sub>T<sub>+</sub>) - (S<sub>+</sub>T<sub>-</sub>) | 0.64 [-0.23, 1.45] | 93.24 | 0.99 |

### IHC staining comparing collagen fiber organization of different samples

Next, we set to find out whether collagen gel combined with spider silk could affect the architecture of collagen matrix after cyclic stretch. We applied Masson’s Trichrome to stain the collagen, whereas blue stain represented the collagen (Figure 3). Results showed that when scaffolds composed of collagen alone were stretched, the gel tended to be thinner, which corresponded to the shrunken shape and reduced stiffness in Figure 3. To the contrary, collagen scaffold containing spider silk showed similar thickness and organized architecture after cyclic stretch. The collagen gels showed reorganization of collagen fibers after 10% cyclic stretching for 24h at 1Hz. Fibers of collagen gel infused with the silk after stretching appeared to be denser and more compact (Figure 3).

**Figure 3.**
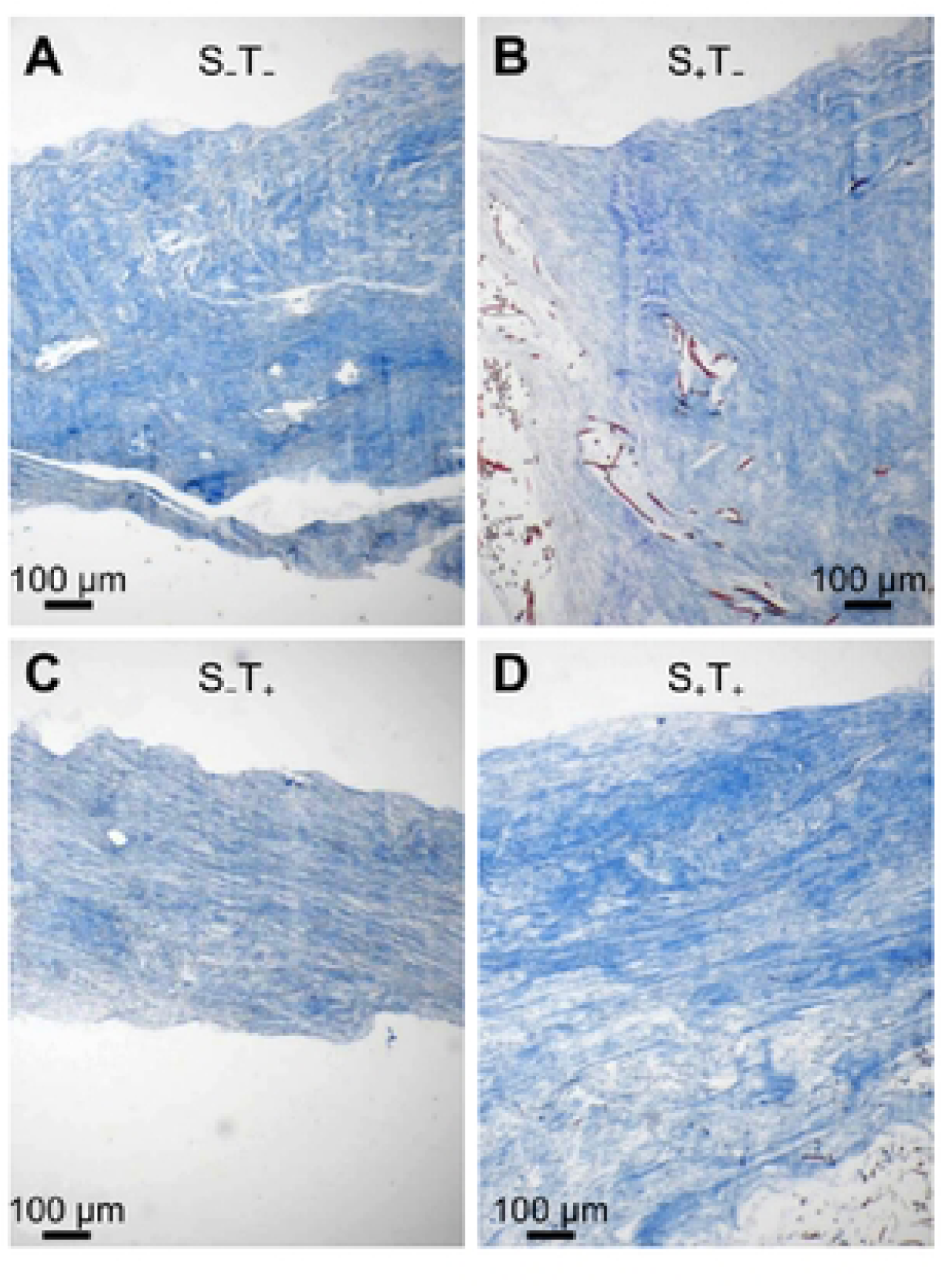
Collagen scaffold containing spider silk showed maintained thickness and organized architecture after cyclic stretch. (A) Gel composed of only collagen without stretching. (B) Gel infused with silk without stretching. (C) Gel composed of only collagen subjected to 10%, 24h cyclic stretch. (D) Gel infused with silk subjected to 10%, 24h cyclic stretch. Abbreviations: S_+_, sample with spider silk; S_−_, sample without spider silk; T_+_, stretched sample; T_−_, unstretched sample.

## Discussion

Silk is a promising biomaterial due to its incredible mechanical properties and biodegradability. On the other hand, collagen gels also show great biocompatibility and biodegradability. Collagen gels exhibit great potential as a matrix for cell cultures, drug delivery carriers, and treatment of burns and sores [36,37,38]. Previously, researchers had attempted to combine recombinant spider silk with collagen, but currently, there are no established techniques for such an attempt. In this study, we decided to use native spider silk collected directly from giant wood spider onto plastic frame to create collagen plus silk construct. We wanted to create a dynamic system composed of collagen and natural spider silk that could be cyclically stretched without losing initial stiffness. Such stretchable scaffold may substitute animal model in future research since untreated collagen readily loses stiffness or becomes shrunken after cyclic stretch [39]. Mechanical stimulation, such as the one we applied in this present study can mimic dynamics of living body [40]. We placed collected natural spider silk onto PDMS membrane and subsequently collagen gel was poured in, and gelation occurred *in situ*. After 10% strain, 24h cyclic 1 Hz stretching was applied to the collagen plus silk construct, we used AFM to confirm the stretch-resistance of the gels. Here we created for the first time a gel construct composed of collagen and natural spider silk. Such novel construct exhibited good stretch-resistant mechanical property as shown in Figure 1 and 2. After being cyclically stretched for 24 hours, our collagen plus silk construct remained intact, was not fragile and had no bacterial growth. Results of AFM measurements showed that the stiffness of such construct was significantly higher than that composed of collagen alone. Additionally, we performed IHC staining to understand the underlying mechanisms of the enhanced mechanical performance. Cyclic stretching would induce the collagen fibers of the gel to reorganize. When the collagen gel was infused with natural spider silk, after cyclic stretching the fibers were reorganized in a more compact, aligned and denser manner.

The microenvironments of cells *in vivo* are actually quite dynamic and they constantly experience various mechanical stimulations such as contraction and relaxation of muscle cells [41], shear stress from blood flow [42], compression in bone tissue and tension in tendons [43]. Recent studies showed that the dynamic interactions between cells and immediate ECM we essential for the survival and growth of cells. Moreover, providing adequate mechanical stimulation may help elicit normal function of cells [44] and might cause cancer cells to die [18]. Berrueta et al. [16] empirically demonstrated that stretching could reduce the size of mouse breast tumor by 52%. Because during early phase of cyclic stretching certain cancer cells would change their structure and size in a unique way, such phenomenon indicates a great applicational potential in early cancer diagnosis [45]. In this study, we demonstrated that infusing collagen gel with natural spider silk could improve mechanical performance of collagen under dynamic conditions. When combined with natural spider silks, collagen gel became more mechanically stable with higher stiffness under cyclic stretching condition, which renders the collagen plus silk construct a good matrix to support cell growth and proliferation while preventing bacterial infiltration [2,3]. This stretchable gel construct composed of natural spider silks may provide an insight into creating matrix with mechanical strength similar to animal tissue. Our next step will be performing cell adhesion experiments and investigating the roles that surface layers of the native silk play in aligning with collagen fibers to find out how spider silk intertwines or disturbs the collagen fibers at microscopic level. We hope that our research will lead to new discoveries of potential application of collagen-based hydrogel in skin wound healing.

Major ampullate spider silk fibers are made up mainly of two proteins – spidroin MaSp1 and MaSp2. A dozen of amino acids were found in the silk, with the dominant ones including glycine, alanine, serine, proline and glutathione tripeptides that were connected with peptide bonds. Spidroins are big (up to 350 kDa) monomers, which core consists of non-repetitive terminal domains with globular structures. Nonrepetitive NTDs and CTDs contribute to fiber formation, while repetitive core is responsible for mechanical strength [46]. Terminal domains are rich in glycine and alanine. Alanine-rich crystalline regions containing anti-parallel β-sheets composed of polyalanine motifs are responsible for tensile strength, while glycine-containing motifs render high extensibility of spider silk [47]. Recent studies showed that GGX motifs might form β-sheets or ordered helical structures, while GPGXX motifs form β-turns. The GPGXX motif forms a spiral that is similar to the beta-turn structure of elastin. The helical structures composed of GGX motifs might play a role in aligning fibers by connecting crystalline beta-sheet regions within the protein and between GGX helices [46]. Collagen, on the other hand, consists of triple helical domains with varying lengths of tri-amino acid repeats Glycine-X–Y. The amino acid composition of collagen is characterized by its high hydroxyproline content. After fibril formation, lysine and proline residues that are hydroxylated by lysyl and prolyl hydroxylase enzymes may stabilize collagen helix formed by three protein monomers named “α chains” [26]. Helical structure of silk fibers and collagen helix might be the key for integration and alignment of those fibers. Crystalline β-sheet structures of spider MA silk could also be involved in collagen peptide linkage. However, the exact mechanism underlying structural integration of collagen and spider silk awaits further studies.

## Acknowledgment

We want to express our sincere gratitude to Ling-Fei Chen for her support with all the documents for the experimental permits and funding.

## Author Contributions

Maryia Tsiareshyna: Data curation; Formal analysis; Investigation; Methodology; Writing -original draft. Sammi Yen-Ting Huang: Visualization; Writing - original draft; Writing - review & editing. Chen-Pan Liao: Statistical analysis; Methodology; Visualization; Writing -original draft. Ming-Jer Tang: Conceptualization; Funding acquisition; Project administration; Resources; Software; Supervision; Tzyy Yue Wong: Data curation; Formal analysis; Investigation; Methodology; Writing -original draft. I-Min Tso: Conceptualization; Funding acquisition; Project administration; Resources; Software; Supervision; Validation; Visualization; Writing - review & editing.

## Competing interesting

The authors declare no conflict of interest.

## Data availability statement

The raw data was public in the following link: https://doi.org/10.6084/m9.figshare.26829685.v2

